# VEGFR2 blockade converts thermally ablative focused ultrasound into a potent driver of T cell-dependent anti-tumor immunity

**DOI:** 10.1101/2025.10.23.683708

**Authors:** Mark R. Schwartz, Nareen Z. Anwar, Lydia E. Kitelinger, David R. Brenin, Patrick M. Dillon, Jonathan R. Lindner, Timothy N.J. Bullock, Matthew R. DeWitt, Richard J. Price

**Affiliations:** Department of Biomedical Engineering, University of Virginia, Charlottesville, VA; Department of Pathology, University of Virginia, Charlottesville, VA; Department of Radiology & Medical Imaging, University of Virginia, Charlottesville, VA; Department of Surgery, University of Virginia, Charlottesville, VA; Department of Medicine, University of Virginia, Charlottesville, VA

## Abstract

Thermally ablative focused ultrasound (TFUS) can induce favorable immune signatures in solid tumors but rarely generates durable systemic immunity or enhances checkpoint inhibition. We tested whether combining TFUS with aVEGFR2, which normalizes tumor vasculature and remodels immune cell composition, triggers T cell–dependent immunity and synergizes with checkpoint inhibition. Subtotal TFUS (∼40% volumetric ablation fraction, matching clinical trial results) was applied to EMT6 tumors in aVEGFR2-treated BALB/c mice. In EMT6 tumors, TFUS or aVEGFR2 alone had no effect on tumor growth, but their combination eradicated 50% of tumors and drove 83% rechallenge rejection, with CD4/CD8 depletion confirming that tumor eradication and rechallenge rejection were T cell–dependent. TFUS + aPD1 yielded modest benefit, whereas triple therapy (TFUS + aVEGFR2 + aPD1) cured 81% of mice and induced durable, T cell–mediated immunity. Thus, VEGFR2 blockade converts clinically relevant TFUS into a potent systemic immunotherapy and, with the addition of checkpoint inhibition, offers a rational approach to combining already-approved therapies for the treatment of solid tumors.

## Introduction

Several minimally and non-invasive ablation approaches, including laser interstitial thermal therapy (LITT), microwave and radiofrequency ablation, cryoablation, irreversible electroporation, and thermal ablation with focused ultrasound (TFUS) ^1^, are clinically deployed to destroy solid tumors. While these ablative techniques are typically leveraged to elicit local cancer cell death as a monotherapy, intratumoral immunomodulation is also known to occur ^2–15^. For example, while TFUS is typically used to achieve coagulative necrosis ^11^, debulk neoplasms ^16^, and temporarily control primary tumor growth ^17–19^, it also induces damage-associated molecular pattern expression and release ^14,20^, and liberates tumor antigen ^21^. Since it is metastatic, not primary, tumor growth and dissemination that causes patient mortality in many cancer indications, it is intriguing to consider how immunological responses to TFUS could be leveraged to drive systemic anti-tumor immunity against metastatic deposits.

Indeed, the so-called abscopal effect has been reported in numerous preclinical small-animal models ^22^ and clinical case reports^23,24^ after primary tumor ablation. However, there is little evidence that TFUS or other ablation modalities, when delivered as local monotherapies, can robustly and reproducibly control distal metastatic disease, despite their ability to modulate the tumor immune landscape. Combining ablation with immune checkpoint inhibitors (ICIs) offers a rational strategy to translate local tumor destruction into systemic, durable immunity. Sizeable subsets of triple-negative breast cancer ^25^, metastatic melanoma ^26^, and pancreatic cancer ^27^ patients are refractory to ICIs, which is primarily due to intratumoral immunosuppression, poorly functioning tumor vessels, and low tumor antigenicity. Combining thermal ablation with ICIs may overcome some of these factors, particularly by temporarily abolishing immunosuppressive intratumoral cells, while releasing tumor antigen ^28^. While preclinical studies show that ablation can potentiate ICIs, clinical results have been inconsistent, with patient response rates frequently below 10% ^29^ when treated with a combination of ablation and ICI ^30^. This underwhelming response is likely due to persistent barriers, such as poor tumor perfusion, hypoxia, and limited immune cell infiltration ^31^. Here, we hypothesize that TFUS could be leveraged to immunologically control distal metastatic disease via logical combinations with approved therapies.

To this end, one potential drug class which may be combined with TFUS to improve patient responses to ICIs is angiogenesis inhibitors, which can reverse immunosuppression in the tumor microenvironment, primarily through modulation of tumor vessel properties ^32–36^. Tumor blood vessel dysfunction is a primary factor in unresponsiveness to anti-cancer drugs, including immunotherapy. This aberrance presents both at the macro-level, with a disorganized vascular network ^37^ that insufficiently perfuses tumor tissue ^38^, and at the micro-level, with an anergic response to inflammatory stimulus and poor capacity for leukocyte extravasation ^39–42^. Low doses of angiogenesis inhibitors have proven effective in temporarily reversing undesirable tumor vessel properties in a process termed “vascular normalization”. This includes improving blood vessel perfusion ^32^, intratumoral hypoxia ^33,34^, adhesion molecule expression ^35,43^, and T cell extravasation into tumors ^41,44^. Although low-dose antiangiogenics alone may not substantially impact tumor growth kinetics, their combination with immunotherapy, such as anti-cancer vaccines ^45–47^, adoptive immune cell transfer ^48–50^, and ICIs ^51–58^, have improved the capacity for immunotherapies to control tumor growth and elicit systemic anti-tumor responses. Since TFUS is also known to elicit anti-tumor immune responses, we expect that the immunostimulatory effects of TFUS may also be enhanced with low-dose angiogenesis blockade.

In these studies, we tested whether these three therapeutic interventions—TFUS, low-dose angiogenesis inhibition, and ICI—can be combined to eradicate solid tumors and generate systemic anti-tumor immunity to resist subsequent rechallenge. To this end, we first developed a partial thermal ablation protocol using TFUS that mimics a clinical trial approach and applied it in multiple syngeneic murine subcutaneous tumor models. After confirming the ability of DC101, a monoclonal antibody that inhibits vascular endothelial growth factor receptor 2 (VEGFR2), to modulate tumor T cell presence without affecting tumor growth, we demonstrated that TFUS combined with DC101 and anti-PD1 (aPD1) robustly controls tumor growth and drives T cell-dependent anti-tumor immunity. These findings position anti-VEGFR2 (aVEGFR2) as a potent enabler of thermal ablation-induced immune responses for treating metastatic disease and highlight the therapeutic potential of integrating energy-based therapies with multimodal-immunomodulation.

## Results

### Influence of TFUS monotherapy on EMT6 tumors

We first developed a TFUS protocol that mimics ablation fractions generated in a recent clinical trial at the University of Virginia (NCT03237572). We employed a FUS system with 4 therapeutic transducers (3.78 MHz) and a central imaging transducer for treatment guidance (Figure 1A), described as System 1 in the Methods section. TFUS was applied at 1.5-mm intervals in the X and Y directions, with treatment planes 2 mm apart (Figure 1B). We confirmed that temperatures >70°C were achieved using thermochromic gels treated with the same parameters (Figure 1B). In addition, we stained ablated EMT6 tumors with 2,3,5-Triphenyltetrazolium chloride (TTC), which reveals metabolically active cells (Figure 1C), to determine the fraction of ablated tumor (Figure 1D). The outer edges of the tumor stained red, the TFUS treated center stained white, and the periablative “transition” zone stained pink, indicating partial TFUS ablation was achieved. Importantly, the fraction of ablated tumor (∼40%) matched that measured in clinical trial NCT03237572 (Figure 1D), wherein ICI was combined with TFUS in late-stage breast cancer patients.

**Figure 1.**
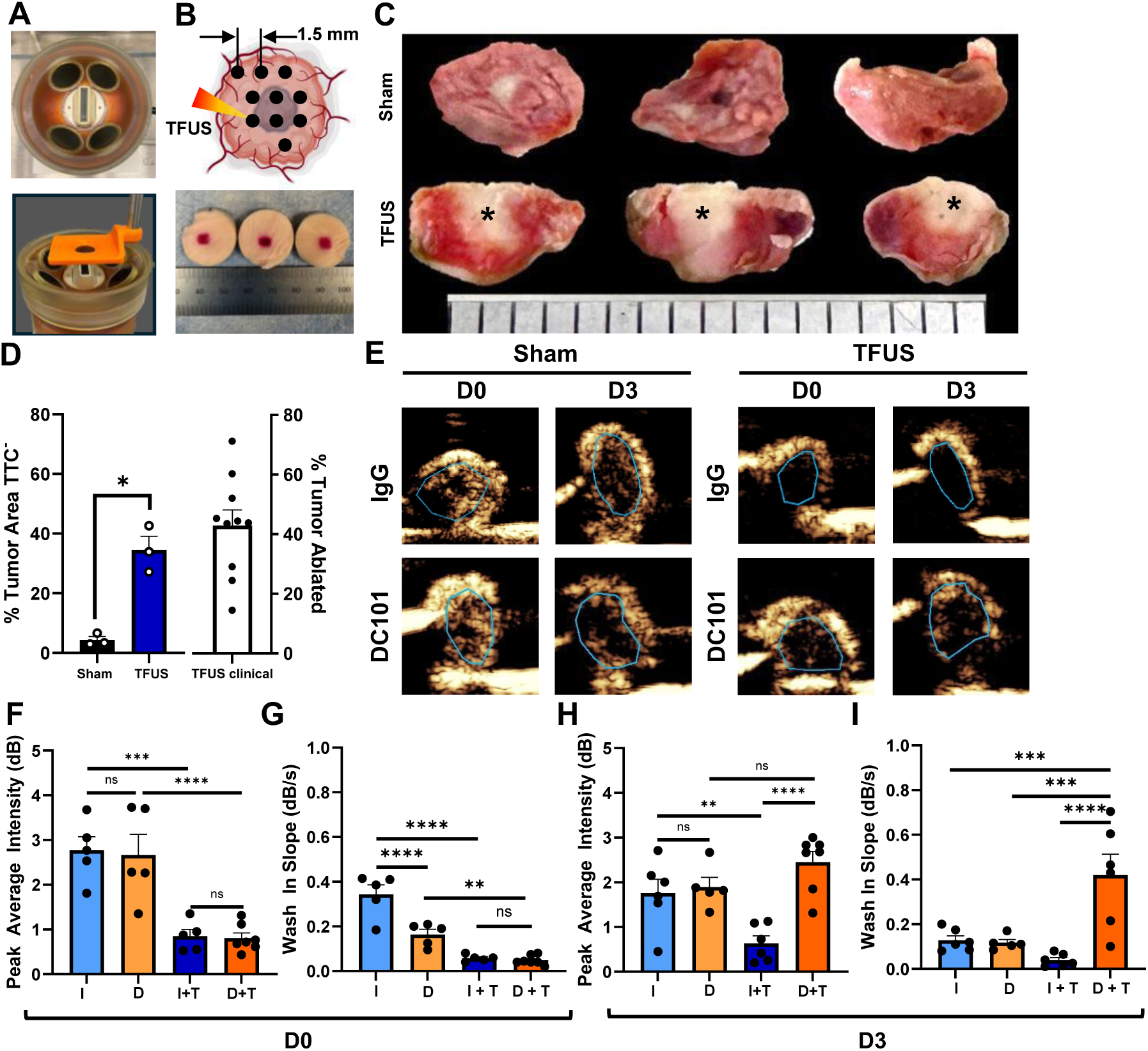
Thermal focused ultrasound equipment and parameters. A) Four-element FUS system with 3.78 MHz transducers. B) Treatments used a 1.5 mm spacing between points and 2 mm between treatment planes. Thermochromic gels with a color change transition to dark red at 70° confirmed temperatures exceeded 70°C when applying FUS at 18 W for 15 s with 1.5 mm spacing. C) Representative TTC staining for TFUS- and Sham-treated EMT6 tumors. Asterisks denote ablated regions. Ruler divisions are 1mm. D) Bar graphs of TTC staining area as a percent of total area, revealing TFUS ablation fraction of EMT6 mouse tumors, and TFUS ablation fractions of primary breast tumors in patients in clinical trial NCT03237572. *P<0.05. Unpaired Welch’s t-test. E) CEUS imaging of EMT6 tumors in IgG- and DC101-treated mice immediately (D0) and 3 days (D3) post-TFUS. Sham-treated mice are shown for comparison. Cyan outlines denote tumor location. F-I) Bar graphs of Peak Average Intensity (F, H) and Wash-in slope metrics (G, I), derived from CEUS imaging, at D0 and D3 for IgG (I) and DC101 (D) treated mice exposed to TFUS (T) ablation or Sham treatment. Two-way ANOVAs with Tukey’s tests. **P<0.01; ***P<0.001; ****P<0.0001.

We next tested how TFUS monotherapy affected the growth of EMT6 tumors, a syngeneic, triple-negative breast cancer model. TFUS application at 14 days post-inoculation (tumor volume of ∼100 mm^3^) was insufficient to control EMT6 tumor growth compared to a sham control (Figure 2A, B). An “area under the curve” metric, which is a measure of integrated tumor burden, yielded an ∼2-fold decrease in tumor burden with TFUS (Figure 2C). However, TFUS improved neither animal survival (Figure 2D) nor tumor eradication (Figure 2E). Thus, this TFUS monotherapy yields modest and transient primary tumor control.

**Figure 2.**
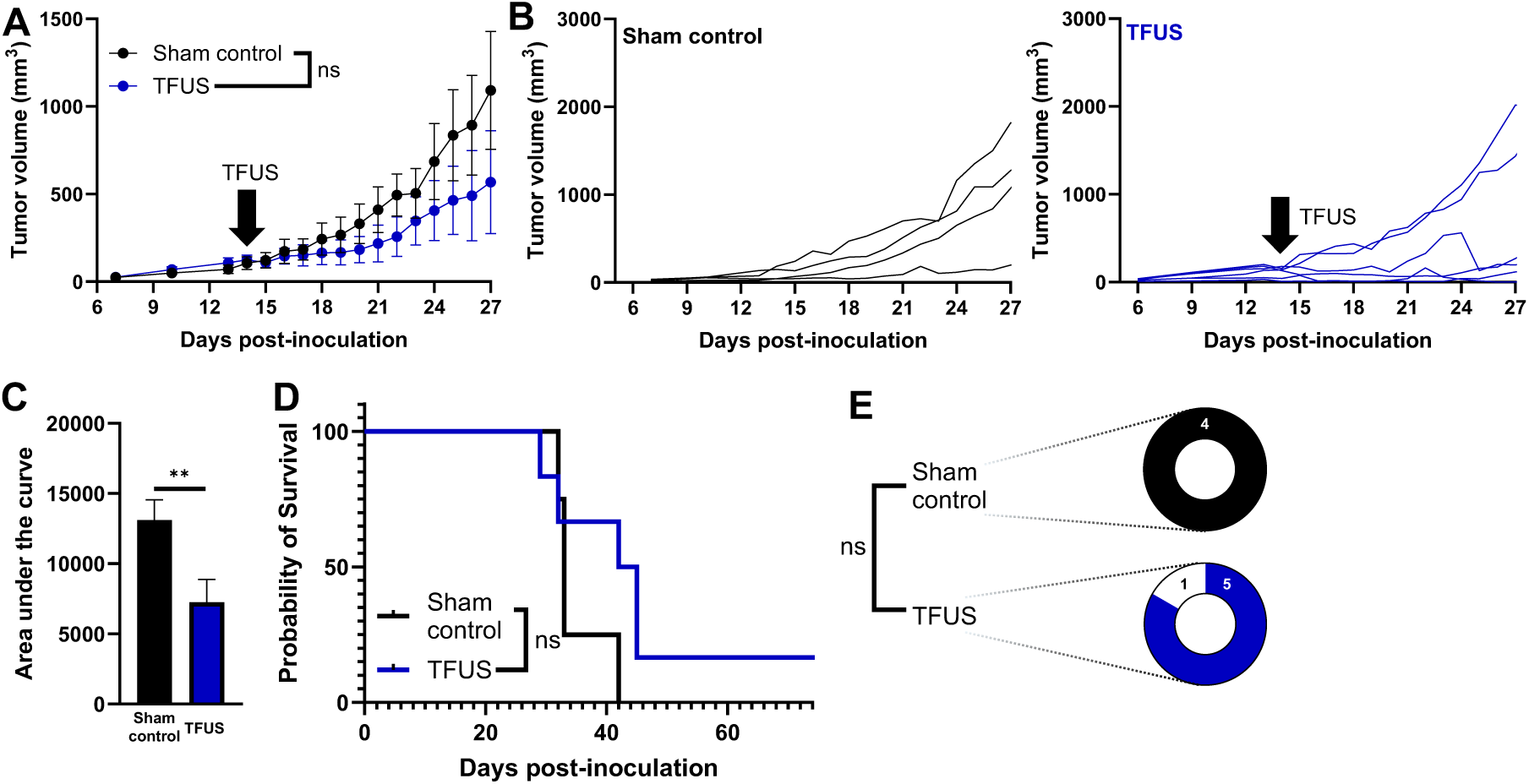
Influence of TFUS monotherapy on EMT6 tumor growth. A) Grouped tumor growth over time. Two-way repeated measures ANOVA. B) Individual growth curves. C) Area under the curve (AUC) for tumor growth. Unpaired Welch’s t-test, **P = 0.0074. D) Kaplan-Meier curve depicting overall survival. Mantel-Cox Log-Rank test. E) Tumor eradication rate. Fisher’s exact test, P = 1.

### VEGFR2 blockade with DC101 beneficially modulates intratumoral T cell composition

We next asked whether aVEGFR2 could synergize with TFUS to control primary tumor growth and drive T-cell dependent anti-tumor immunity. We first tested whether aVEGFR2 beneficially remodels the immune landscape in EMT6 tumors. DC101 was administered i.p. and EMT6 tumor growth was tracked beginning 7 days after inoculation (Figure 3A). A subset of tumors was harvested 14 days post-inoculation when tumors measured ∼100 mm^3^ and assessed for T cell representation by flow cytometry, using the gating strategy in Figure S1. The number and share of CD8^+^ T cells in these tumors were unchanged by DC101 (Figure 3B, C). However, the number and share of CD4^+^Foxp3^+^ regulatory T cells (T_reg_) decreased by ∼3-fold and ∼1.5-fold, respectively (Figure 3D, E). As a result, the CD8/Treg ratio, a predictor of improved clinical outcomes ^59,60^, nearly doubled with DC101 treatment (Figure 3F). Additionally, about half of the CD8^+^ cells were PD1^+^ following either IgG or DC101 treatment (Figure 3G). However, DC101 monotherapy did not control tumor (Figure 3H, I), improve integrated tumor burden (Figure 3J), extend animal survival (Figure 3K), or eradicate tumors (Figure 3L). Therefore, DC101 monotherapy induces favorable immune changes but fails to impact tumor growth or eradication.

**Figure 3.**
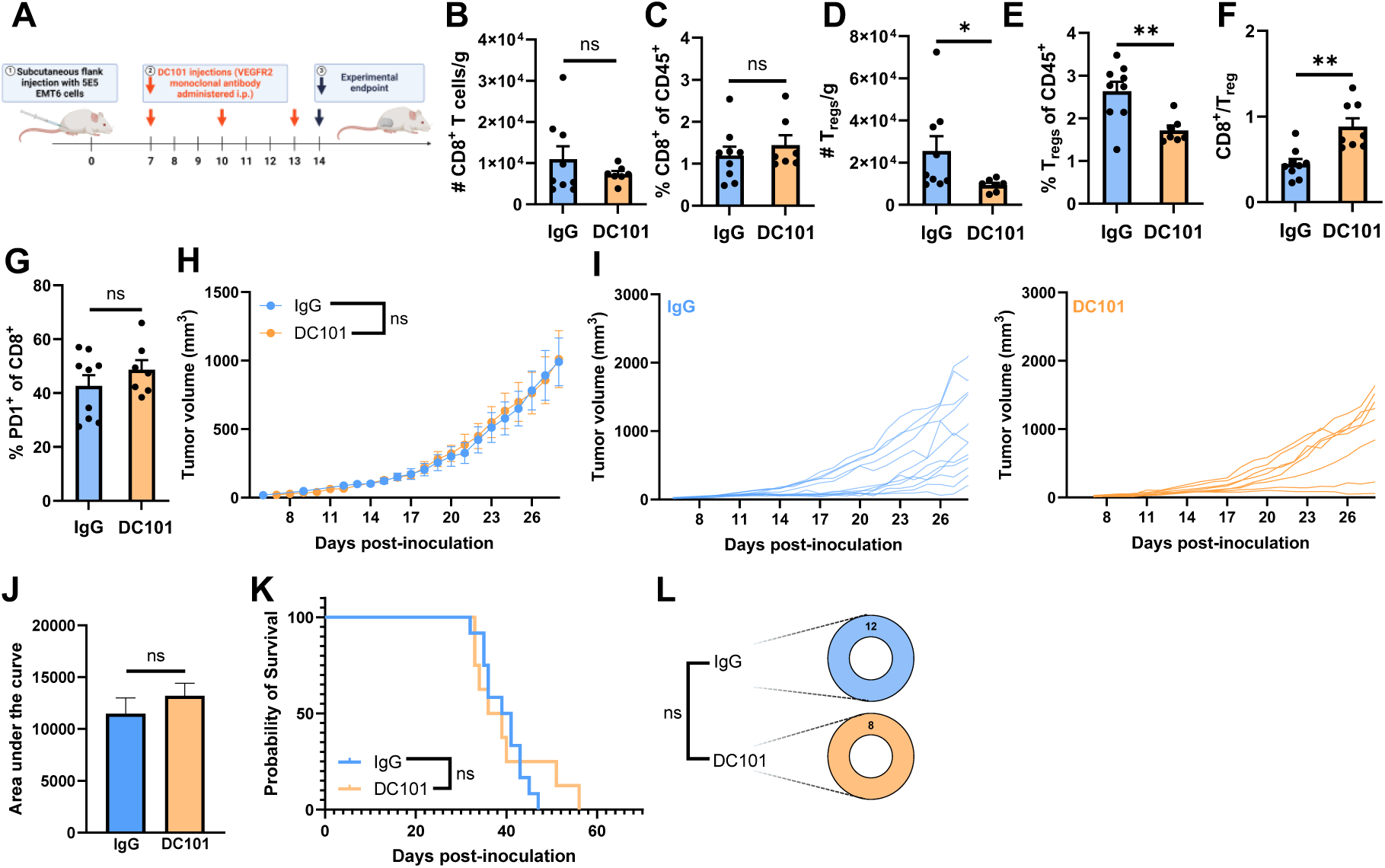
DC101 modulates T cell presence without controlling tumor growth. A) Timeline for inoculation and treatment. B-F) Bar graphs of CD8^+^ T cells per gram tumor (B), percentage of CD45^+^ cells that are CD8^+^ T cells (C), T_reg_ per gram tumor (D), percentage of CD45^+^ cells that are T_reg_ (E), and ratio of CD8^+^ T cells to T_reg_ (F). Welch’s t-tests. *P <0.05; **P<0.01. G) Percentage of CD8^+^ T cells that are also PD1^+^. Welch’s t-test. H) Grouped tumor growth over time. Two-way repeated measures ANOVA. I) Individual growth curves. J) Area under the curve (AUC) for tumor growth. Welch’s t-test. K) Kaplan-Meier curve depicting overall survival. Mantel-Cox log-rank test. L) Tumor eradication fraction. Fisher’s exact test, P = 1.

### VEGFR2 blockade with DC101 accelerates post-ablation perfusion recovery

Because VEGFR2 blockade is known to enhance tumor perfusion, we hypothesized that perfusion recovery after TFUS would be accelerated by DC101. To test this, we assessed tumor perfusion with contrast-enhanced ultrasound imaging (CEUS) (Figure 1E). Bulk tumor perfusion was assessed using peak average intensity and wash-in slope metrics, consistent with perfusion analysis of tumor vasculature. Notably, both peak intensity and wash-in slope were decreased between 70 and 80% in control and DC101 groups immediately post-TFUS (D0), applied with FUS System 1, with no difference in peak intensity between IgG and DC101. Moderate differences in wash-in slope were observed and likely explained by vascular normalization effects on blood flow velocity, but not total perfused volume (Figure 1F, G). At 3 days post-ablation, both perfusion metrics were still 65-75% lower in IgG controls treated with TFUS. However, in contrast, ablated tumors in DC101-treated mice exhibited complete recovery of both perfusion metrics by 3 days after TFUS (Figure 1H, I).

### DC101 synergizes with TFUS to control tumor growth, extend survival, and induce anti-tumor immunity

After establishing that aVEGFR2 remodels tumor immune landscape and improves post-TFUS perfusion recovery, we tested whether it synergizes with TFUS in the treatment of EMT6 tumors (Figure 4A). Here, DC101 + TFUS, applied with FUS System 1, significantly controlled EMT6 tumor growth when compared to DC101 and IgG control groups (Figure 4B, C). At 2 weeks post-TFUS, the DC101 + TFUS-treated tumor volume was fully 5-fold lower than IgG control and 3.5-fold lower than TFUS monotherapy, constituting a synergistic relationship between DC101 and TFUS (Figure 4D). as assessed by both the response additivity and Bliss independence models^61^. The response additivity model defines a relationship as synergistic when the effect from two distinct therapies is greater than the sum of the individual therapies’ effects, while the Bliss independence model finds synergy when the effect of two therapies acts on different sites of action and is greater than the sum and the product of the two therapies effects. We employ both models to set a higher synergy threshold. Tumor control was reflected in the AUC metric as well, with a 3.5-fold decrease in tumor burden for DC101 + TFUS compared to IgG control and a 2-fold decrease compared to TFUS (Figure 4E). Survival was also extended significantly in the DC101 + TFUS group (Figure 4F) and 50% of tumors treated with DC101 + TFUS were entirely eradicated, an improvement from 17% of tumors eradicated by TFUS (Figure 4G). Neither the DC101 group nor the IgG group yielded tumor eradication (Figure 4G). Similar results were observed in the syngeneic, triple negative breast cancer model, 4T1, where we observed a significant growth control benefit with DC101 + TFUS (Figure S2) when TFUS was applied using FUS System 2. A trending decrease in volume of lung metastases was also observed at 4 weeks post-inoculation (Figure S2).

**Figure 4.**
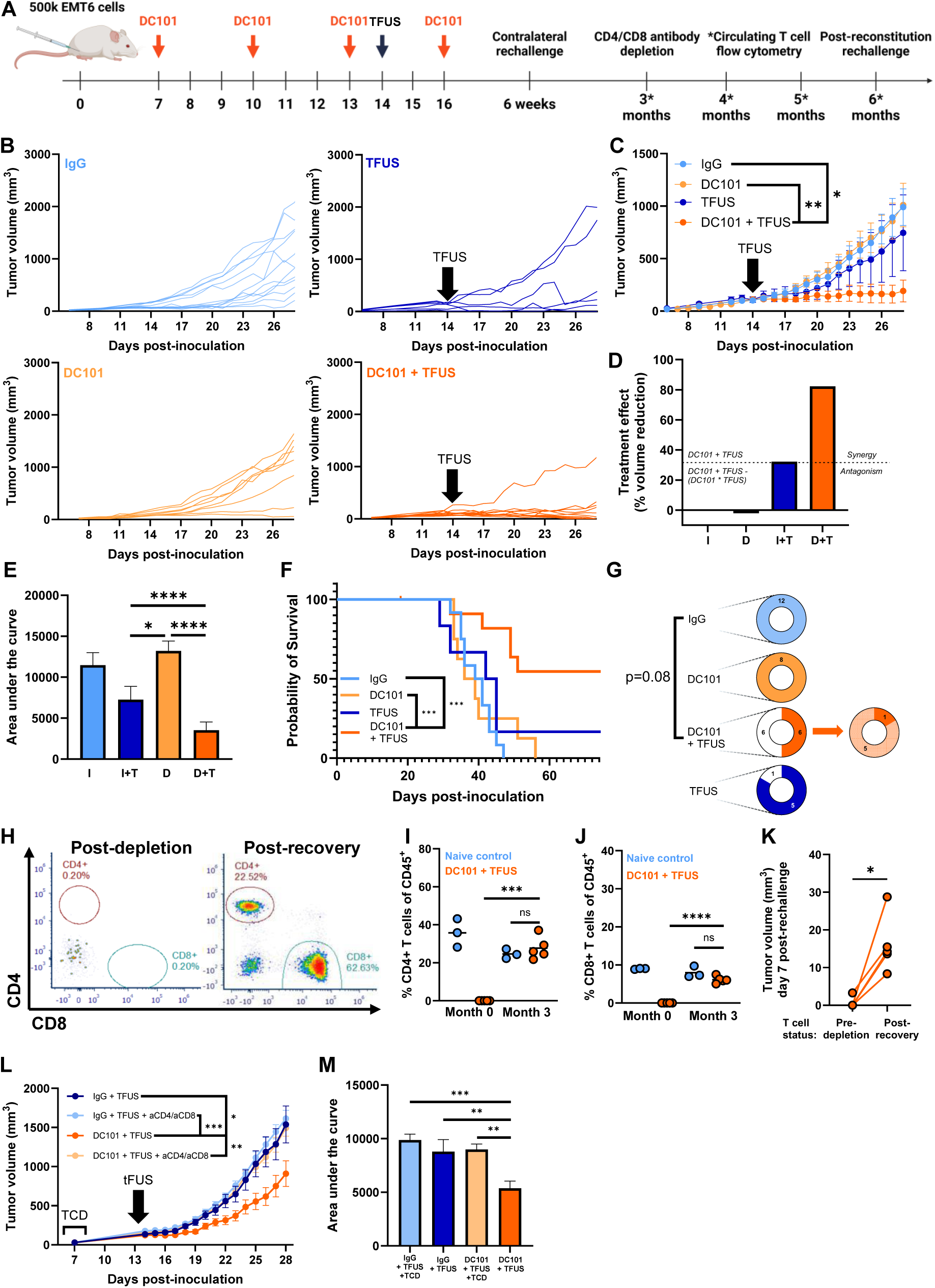
TFUS synergizes with DC101 to control primary tumor growth and generate a systemic anti-tumor response. A) Timeline for inoculation and treatment. B) Individual growth curves. C) Grouped tumor growth over time. Two-way repeated measures ANOVA; *P = 0.0207, **P = 0.007. D) Graphical representation of the combinatory relationship between DC101 (D) and TFUS (T). I = IgG control. E) Area under the curve (AUC) for tumor growth. D = DC101; T = TFUS; I = IgG control. Two-way ANOVA with Tukey’s test. ****P < 0.0001, *P = 0.0428. F) Kaplan-Meier curve depicting overall survival. Mantel-Cox log-rank test. ***P < 0.001. G) Tumor eradication (white shading) and rechallenge rejection (light orange shading) fractions. Fisher’s exact test. H) Flow cytometry scatter plots of circulating CD4 and CD8 T cells immediately post-depletion and post-recovery 3 months later. I) Percentage of CD45^+^ cells in circulation that are CD4 T cells, immediately post-depletion and 3 months later. Two-way ANOVA with Tukey’s test. ****P < 0.0001. J) Percentage of CD45^+^ cells in circulation that are CD8 T cells, immediately post-depletion and 3 months later. Two-way ANOVA with Tukey’s test. ****P < 0.0001. K) Tumor volume, 7 days post-rechallenge, in the same mice, pre-T cell depletion and post-T cell recovery. Paired t-test. *P = 0.011. L) Grouped tumor growth for TFUS ablation, with and without DC101 pretreatment, after CD4 and CD8 T cell depletion (TCD) or control treatment. Two-way, repeated measures ANOVA. *P = 0.0112, ***P = 0.0007, **P = 0.0090. M) Area under the curve (AUC) for tumor growth. Two-way ANOVA with Tukey test. ***P < 0.001, **P < 0.01.

To then test whether DC101 + TFUS-treated mice with eradicated primary tumors harbored systemic immunity against EMT6 tumors, we performed EMT6 rechallenges in the contralateral flank. Here, 83% of mice rejected the rechallenge (Figure 4G), suggesting the presence of systemic anti-tumor immunity. To verify that tumor rejection was T cell-dependent, we then depleted CD4 and CD8 T cells, followed by a second rechallenge. Three months after CD4/CD8 depletion, T cell reconstitution was complete (Figure 4H-J). Post-recovery, EMT6 tumors grew significantly larger than they did in the pre-depletion rechallenge experiment (Figure 4K), indicating that the control of contralateral tumors conferred by the combination treatment was T cell dependent. After this time point, most tumors rechallenged post-reconstitution did regress, which is consistent with the presence of tissue resident memory T cells in the skin and lungs that are difficult to deplete with systemic αCD4 and αCD8 antibodies ^62–64^.

We also tested whether T cells were necessary for primary tumor control. CD4/CD8 depletion (TCD) prior to ablation revealed that T cells are responsible for primary tumor growth control with DC101 + TFUS treatment, but not with TFUS alone (Figure 4L, M). Together, these data illustrate that DC101 and TFUS synergize to control primary tumor growth and generate long-term immunity, both of which depend on T cells.

### aPD1 combined with TFUS additively controls tumor growth and augments anti-tumor immunity

Since vascular and immune remodeling achieved with DC101 cooperated with TFUS, we asked whether another well-studied immune remodeling strategy, aPD1, also cooperates with TFUS. EMT6 tumors are partially responsive to similar drugs ^65–67^, mimicking the clinical response of TNBC patients to ICIs^68^. We dosed EMT6-bearing mice with aPD1 (4 doses using a standard concentration and timing for mice) and applied TFUS with FUS System 1 on day 14 (Figure 5A). We observed a trending decrease (p=0.064) in tumor volume for the aPD1 + TFUS group when compared to IgG (Figure 5B, C) and no significant differences when compared to TFUS and PD1 monotherapies, demonstrating a cooperative relationship between both treatments (Figure 5D). The AUC metric revealed an ∼3-fold decrease in integrated tumor burden with aPD1 + TFUS compared to IgG control (Figure 5E), as well as a trending (p=0.12) survival advantage compared to IgG control (Figure 5F). Notably, this protocol was repeated in a syngeneic melanoma tumor model, YUMMER1.7, wherein similar results were observed (Figure S3). ∼55% of YUMMER1.7 tumor-bearing mice treated with aPD1 + TFUS exhibited tumor eradication (Figure 5F) as compared to only ∼15% of aPD1-treated mice.

**Figure 5.**
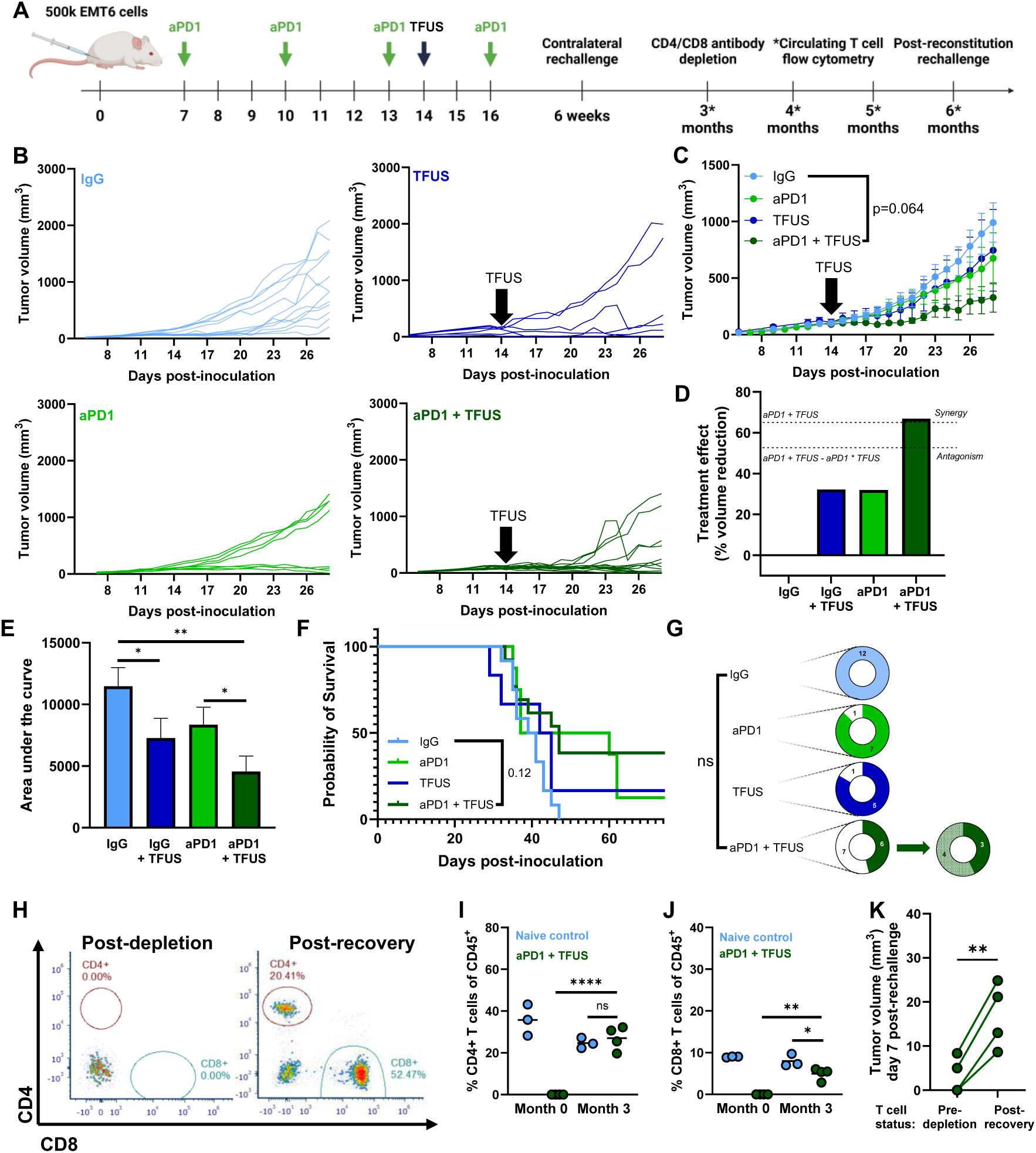
TFUS additively cooperates with aPD1 to control primary tumor growth and induce anti-tumor immunity. A) Timeline for inoculation and treatment. B) Individual growth curves. C) Grouped tumor growth over time. Two-way repeated measures ANOVA. D) Graphical representation of the combinatory relationship between aPD1 and TFUS. E) Area under the curve (AUC) for tumor growth. Two-way ANOVA with Tukey’s test. *P <0.05; **P = 0.0011. F) Kaplan-Meier curve depicting overall survival. Mantel-Cox log-rank test. G) Tumor eradication (white shading) and rechallenge rejection (light green shading) fractions. Fisher’s exact test. P = 0.5. H) Flow cytometry scatter plots of circulating CD4 and CD8 T cells immediately post-depletion and post-recovery 3 months later. I) Percentage of circulating CD45^+^ cells that are CD4 T cells, immediately post-depletion and 3 months later. Two-way ANOVA with Tukey’s test. ****P < 0.0001. J) Percentage of circulating CD45^+^ cells that are CD8 T cells, immediately post-depletion and 3 months later. Two-way ANOVA with Tukey’s test. *P<0.05; **P <0.01. K) Tumor volume, 7 days post-rechallenge, in the same mice, pre-T cell depletion and post-T cell recovery. Paired t-test. **P < 0.01.

To mirror the DC101 + TFUS experiment, mice with eradicated tumors were rechallenged contralaterally, with ∼60% of rechallenges rejected (Figure 5G). After depleting CD4 and CD8 T cells and allowing circulating T cells to repopulate (Figure 5H-J), mice were again rechallenged contralaterally. Tumors grew faster post-reconstitution than they did pre-depletion (Figure 5K). Again, this suggests T cell memory dependency on the anti-tumor immune response was generated in most of the tumor-eradicated mice.

### DC101 and aPD1, combined with TFUS, robustly controls EMT6 tumor growth, confers survival advantage, and induces anti-tumor immunity

DC101 and aPD1 were individually capable of improving the anti-tumor effects of TFUS. Additionally, anti-angiogenics like DC101 enhance anti-tumor immunity induced by ICIs like aPD1^51–58^. As such, we tested whether combining these three therapeutics would induce more potent tumor control than TFUS or DC101 + aPD1 alone. We administered 4 doses of DC101 and aPD1 and applied TFUS using FUS System 1 on day 14 (Figure 6A). The DC101 + aPD1 + TFUS-treatment yielded exceptional primary tumor control (Figure 6B, C), representing a synergistic relationship (Figure 6D). The triple combination reduced AUC 5-fold compared to IgG control and 3-fold compared to TFUS, with a 3-fold decrease compared to DC101 + αPD1 (Figure 6E). The triple combination therapy also conferred a significant survival benefit (Figure 6F). DC101 + αPD1 eradicated ∼30% of the tumors, while DC101 + αPD1 + TFUS eradicated ∼80% of the tumors, which constitutes a significant eradication benefit compared to IgG control and a strong trend (p=0.057) toward a benefit compared to TFUS (Figure 6G).

**Figure 6.**
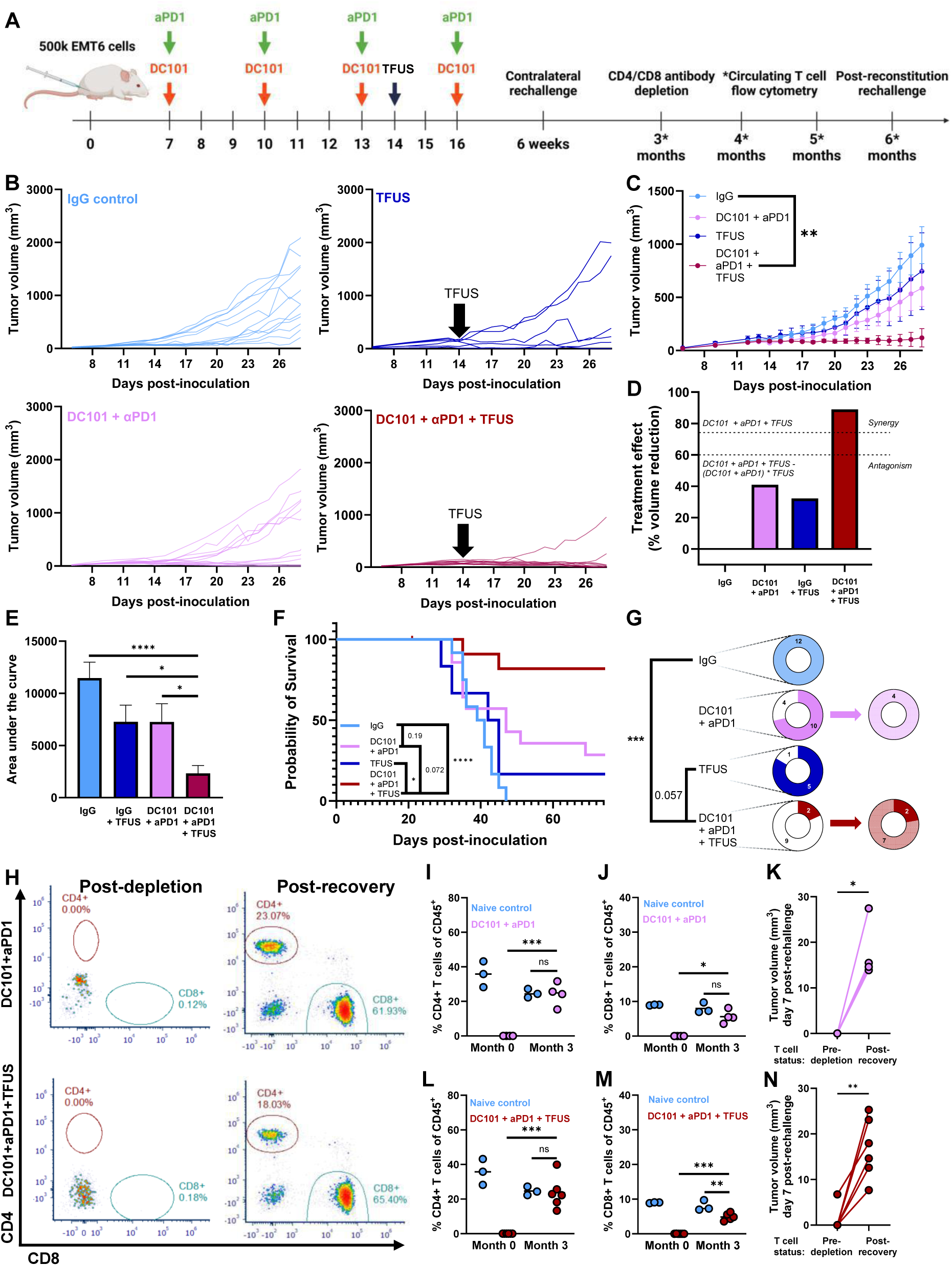
TFUS, DC101, and aPD1 potently cooperate to eradicate tumors and generate a systemic anti-tumor response. A) Timeline for inoculation and treatment. B) Individual growth curves. C) Grouped tumor growth over time. Two-way repeated measures ANOVA. **P = 0.006. D) Graphical representation of the combinatory relationship between DC101 + aPD1 and TFUS. E) Area under the curve (AUC) for tumor growth. Two-way ANOVA with Tukey’s test. ****P < 0.001, *P <0.05. F) Kaplan-Meier curve depicting overall survival. Mantel-Cox log-rank test. ****P < 0.0001, *P = 0.0384. G) Tumor eradication (white shading) and rechallenge rejection (pink shading; light red shading) fractions. Fisher’s exact test. ***P = 0.0006. H) Flow cytometry scatter plots of circulating CD4 and CD8 T cells immediately post-depletion and post-recovery 3 months later. I) Percentage of circulating CD45+ cells that are CD4 T cells, immediately post-depletion and 3 months later. Two-way ANOVA with Tukey’s test. ***P < 0.001. J) Percentage of circulating CD45+ cells that are CD8 T cells, immediately post-depletion and 3 months later. Two-way ANOVA with Tukey’s test. *P <0.05. K) Tumor volume, 7 days post-rechallenge, in the same mice, pre-T cell depletion and post-T cell recovery. Paired t-test. *P = 0.0115. L) Percentage of circulating CD45^+^ cells that are CD4 T cells, immediately post-depletion and 3 months later. Two-way ANOVA with Tukey’s test. ***P < 0.001. M) Percentage of circulating CD45+ cells that are CD8 T cells, immediately post-depletion and 3 months later. Two-way ANOVA with Tukey’s test. **P <0.01; ***P <0.001. N) Tumor volume, 7 days post-rechallenge, in the same mice, pre-T cell depletion and post-T cell recovery. Paired t-test. **P = 0.0026.

Similar to previous experiments, we contralaterally rechallenged the mice with eradicated tumors. All DC101 + αPD1 mice rejected their rechallenge, with ∼75% of the DC101 + αPD1 + TFUS mice rejecting rechallenge (Figure 6G). Following CD4 and CD8 T cell depletion and reconstitution (Figure 6I-J, L-M), mice were contralaterally rechallenged again. Here, we observed faster tumor growth post-reconstitution than we did in the same mice before CD4/CD8 T cell depletion (Figure 6K, N), indicating the rechallenge rejection observed following primary tumor eradication was T cell-dependent.

## Discussion

We aimed to develop a pharmacological strategy that converts TFUS into a non-invasive approach for generating T cell-dependent immunity against solid tumors. Working in a mouse EMT6 tumor model, we first established a partial TFUS regimen that mimicked the average volumetric ablation percentage (∼40%) measured in a clinical trial evaluating TFUS + aPD1 combination approaches in late-stage breast cancer patients (NCT03237572). At this ablation percentage and as a monotherapy, TFUS temporarily slowed tumor growth but did not enhance long-term outcomes, recapitulating typical responses of solid tumors to TFUS. We then leveraged the ability of VEGFR2 blockade to ameliorate intratumoral T cell composition and improve tumor perfusion following ablation to convert partial TFUS into a curative treatment for a substantial fraction of tumor-bearing mice. Rechallenge and CD4/CD8 T cell depletion experiments confirmed the generation of T cell-dependent anti-tumor immunity. Because ∼1/2 of the CD8 effector T cells in EMT6 tumors were PD1^+^, we also evaluated TFUS in combination with aPD1. This combination was only moderately effective, failing to match the DC101 + TFUS combination’s ability to control and eradicate tumors. Finally, combining DC101, aPD1, and TFUS eradicated nearly all primary tumors and, once again, resulted in robust T cell-dependent anti-tumor immunity. Together, these results identify VEGFR2 inhibition as a therapeutic toggle for TFUS, capable of switching it from a tumor debulking modality into a potent driver of T cell-dependent anti-tumor immunity, with further efficacy conferred by the addition of aPD1. Given that both the TFUS ablation scheme and aPD1 are clinically approved, the results support rapid translation of this combinatorial approach to a clinical trial.

Similar to the findings we present, other studies have combined TFUS with immunotherapies to control tumor growth^17,19,69–82^. One key variable in such studies is TFUS ablation fraction, which was set to ∼40% in our study (Figure 1D). We chose this value to match the TFUS ablation fraction applied to late-stage breast cancer patients in clinical trial NCT03237572 (Figure 1D), wherein augmented T cell staining has been observed in peri-ablative regions. Thus, the choice of ablation fraction enhanced the translational potential of the current study. The choice of partial ablation over complete ablation is also supported by other factors. A comparison between partial and total ablation of both MC38 and B16 tumors found that partially ablated tumors experienced greater dendritic cell infiltration and a greater share of mature, intratumoral dendritic cells 1 day post-TFUS^83^. Yet another factor is the potential for partial ablation to increase drug delivery. For example, when partial TFUS is applied as a single point or circular pattern to solid tumors^70,80^, yielding lesions that are generally comparable to those seen in our tumors post-ablation, the delivery of small molecules (gadoteridol), albumin, and liposomes is enhanced^70,80^. In a similar vein, we have shown that nanoliposome delivery is increased 24 hours post-partial ablation of 4T1 tumors^84^. Thus, in the current study, one possible mechanism of enhanced therapeutic benefit with DC101 is the further augmentation of post-ablation drug delivery due to improved tumor perfusion recovery after TFUS (Figure 1F-I). This putative mechanism could also partially explain our finding that, while aPD1 + TFUS yielded only modest cooperation (Figure 5), adding DC101 greatly enhanced primary tumor growth control (Figure 6).

Another key variable is the timing of drug administration relative to ablation. Several groups report success in priming tumors with drugs like TLR agonists and ICIs neoadjuvant to partial ablation^17,19,69–72,75,78,80^. For example, administering TLR agonists and ICIs neoadjuvant to TFUS, as opposed to adjuvant to TFUS, was found to be necessary to control distal tumor growth and promote CD8 T cell trafficking into distal tumors^70^. Here, we took a similar approach and administered 3 neoadjuvant doses of VEGFR2 blockade before TFUS. To ensure that VEGFR2 blockade was impacting EMT6 tumors, we used CD8/T_reg_ ratio as a biomarker to verify beneficial remodeling of tumor immune landscape before TFUS treatment (Figure 3F).

In addition to the novel deployment of aVEGFR2 in combination with TFUS and aPD1 to achieve complete eradication of most (81%) treated primary tumors (Figure 6), we believe one particularly unique and powerful aspect of our study is the mechanistic confirmation that sustained anti-tumor immunity after primary tumor eradication was T cell-dependent. This was evidenced by initial rechallenge rejection in most mice with primary tumor eradication, followed by subsequent rechallenge acceptance in all mice after they had undergone both complete CD4/CD8 T cell depletion from blood and full recovery of circulating CD4/CD8 populations. Other studies have shown that combining partial TFUS with immunotherapies generates primary tumor control, as well as distal tumor control via the abscopal effect^69,70,85^. Such studies also often show immune cell infiltration into these distal tumors, providing evidence for immunological control. Moreover, similar to our studies, contralateral rechallenge approaches have also been used to infer the generation of systemic anti-tumor immunity^83^. Nonetheless, to our knowledge, none of these studies combining partial TFUS with other drugs have deployed T cell depletion and recovery to mechanistically verify that contralateral tumor control or rechallenge rejection is indeed T cell-dependent. This constitutes a long-standing and significant knowledge gap in the field that our study now addresses. Notably, we also confirmed that primary tumor control with aVEGFR2 in combination with TFUS was also CD4/CD8 T cell dependent, and it is reasonable to postulate that CD4/CD8 recruitment to the primary tumor from the bloodstream was boosted by improved tumor perfusion recovery after TFUS in DC101-treated mice. We also note that the finding of T cell-dependent primary tumor control mirrors a previous study from our group wherein we deployed both Rag1^-/-^ knockout mice and CD4/CD8 T cell depletion to verify that primary tumor control with TFUS in combination with gemcitabine^19^ is T cell dependent.

Aside from TFUS, other ablative FUS modalities have also been combined with drug administration to drive systemic anti-tumor immunity, and it is informative to consider our results in light of these other studies. One such modality is histotripsy, which mechanically fractionates tumor tissue with minimal heating and has been reported to be more effective than TFUS in enhancing ICIs in certain preclinical cancer models^86^. Histotripsy activates dendritic cells and promotes tumor antigen acquisition^87^, and it has been argued that the lack of tissue heating following histotripsy may preserve more antigen intact than TFUS regimens which could denature antigen. While these studies highlight the promise of histotripsy as an immunomodulatory ablation modality, our results indicate that partial tumor ablation with TFUS still retains significant promise as a means for augmenting immunotherapy and driving systemic anti-tumor immunity.

## Methods

### Cell and animal maintenance

The EMT6 and 4T1 cell lines were purchased from ATCC. The YUMMER1.7 cell line was gifted by the laboratory of Timothy N.J. Bullock, who was provided with the cells by the laboratory of Marcus Bosenberg. EMT6 cells were maintained in 1X DMEM + 4.5 g/L D-glucose + L-Glutamine (Gibco #11965-092) supplemented with 10% Fetal Bovine Serum (FBS, Gibco #16000-044). 4T1 cells were maintained in 1X RPMI 1640 + L-Glutamine (Gibco #11875-093) supplemented with 10% Fetal Bovine Serum (FBS, Gibco #16000-044). YUMMER1.7 cells were maintained in 1X DMEM/F12 (1:1) + L-Glutamine + 15 mM HEPES supplemented with 10% Fetal Bovine Serum (FBS, Gibco #16000-044) and 1% MEM Non-Essential Amino Acids (NEAA, Gibco # 11140-050). All cells were maintained in culture at 37°C and 5% CO_2_ (Thermo Fisher Scientific, Heracell 150i Cat#51-032-871). Thawed cells were maintained in logarithmic growth phase for all experiments, did not exceed 12 passages from the time of purchase, and tested negative for mycoplasma prior to freezing.

All animal experiments adhered to ethical guidelines and regulations approved by the University of Virginia Animal Care and Use Committee. The animals were housed in accordance with standard laboratory conditions, maintaining a temperature of 22°C and a 12-hour light/12-hour dark cycle and supplied food ad libitum. 7-10 week-old female BALB/c or C57BL/6 mice were purchased from Jackson Laboratories (Jax #000651) and acclimated for at least 48 hours in our animals facilities. To prepare the animals for inoculations, they were anesthetized with ketamine/dextomitor cocktail. Their right flanks were shaved and 5×10^5^ EMT6 cells, 4×10^5^ 4T1 cells, or 3×10^5^ YUMMER1.7 cells were subcutaneously injected in 100 µL 1X PBS (Gibco #10010-023) with a 25G x 1 ½ inch needle (BD PrecisionGlide Needle #305127) into the right flank of the mice, allowed to rest for 30 minutes, and woken up with antisedan. Tumor outgrowth was assessed with digital calipers, with tumor volume = (length x width^2^/2). Seven days following inoculation, mice were randomly placed in experimental groups while matching the starting tumor mean volume and minimizing intragroup variation. When appropriate, animals were rechallenged with inoculations following the same procedure, but on the left flank.

### DC101 therapy

For VEGFR2 blockade, mice were injected with 5 or 10 mg/kg DC101 (DC101 #BE0060 BioXCell) or appropriate IgG antibody control (rat IgG1 HRPN #BE0088 BioXCell) diluted in sterile 1X PBS (Gibco #10010-023). DC101 and IgG were prepared the day of injections, and the 5 or 10 mg/kg calculation was determined at the day 7 size matching and continued throughout the duration of the experiment. Mice were interperitoneally injected with 100 µL of diluted DC101 or IgG at starting on day 7 post-inoculation, with 2 or 3 additional doses administered 3 days apart.

### PD-1 blockade therapy

For PD-1 blockade, mice were injected with 200 μg aPD1 (RMP1-14 #BE0146 BioXCell) or appropriate IgG antibody control (rat IgG2a 2A3 #BE0089 BioXCell) diluted in sterile 1X PBS (Gibco #10010-023). DC101 and IgG were prepared the day of injections, and the 10 mg/kg calculation was determined at the day 7 size matching and continued throughout the duration of the experiment. Mice were interperitoneally injected with 100 µL of diluted aPD1 or IgG at starting on day 7 post-inoculation, with 2 or 3 additional doses administered 3 days apart.

### T cell depletions

For T cell depletions, anti-CD8 (2.43 clone; Bio X Cell) and anti-CD4 (GK1.5 clone; Bio X Cell) were diluted in sterile 1X PBS (Gibco #10010-023) and administered intraperitoneally daily for three days. Mice were injected with 100 µg of each antibody on each of these three days.

### In vivo ultrasound-guided partial thermal ablation

System 1: This system consists of four 3.78 MHz single-element transducers (SU-102, Sonic Concepts), each of 33 mm diameter and 55 mm radius of curvature, with a 3.78 MHz center frequency. These four transducers are embedded in a solid resin and are confocally aligned for a single active aperture of 66 mm. The system is powered by a 200W acoustic amplifier (Electronics & Innovation 1020L) driven by an arbitrary function generator (Tektronix AFG3022C) registered to an ultrasound imaging transducer (MS200, center frequency 30 MHz, FUJIFILM Visualsonics). A degassed water bath at 37°C acoustically coupled mice to both the therapeutic and imaging ultrasound transducers. After acoustic coupling, tumor positioning was driven by a motorized 3D motion stage and the tumor was identified with B-mode ultrasound imaging. Tumors were treated with the transducer operated in continuous wave mode at 18W power for 15 seconds per point, with each treatment point 1.5 mm apart. Tumors were treated in 2 or 3 planes of sonication, each 2 mm apart.

System 2: This system consists of a 64 mm single-element 3.3 MHz transducer (Sonic Concepts) powered by a 400 W amplifier (E&I) orthogonally registered to an 8 MHz linear ultrasound imaging array (Siemens). A degassed water bath at 37°C acoustically coupled mice to both the therapeutic and imaging ultrasound transducers. After acoustic coupling, tumor positioning was driven by a motorized 3D motion stage and the tumor was identified with B-mode ultrasound imaging. A grid of points 3 mm apart was overlayed onto the B-mode images. Tumors were treated with the transducer operated at 3MHz in continuous wave mode at 15W power for 10 seconds per point, and were treating in two planes of sonication, each 2 mm apart.

### Contrast-enhanced ultrasound imaging (CEUS)

To prepare the animals for CEUS imaging, they were anesthetized with a ketamine/dextomitor cocktail. The right flanks were then shaved. Mice were catheterized in their tail vein, then placed into System 1 for imaging. B-mode imaging was used to position the imaging transducer for CEUS image acquisition. Mice were injected with 40 uL of neutral lipid microbubbles at a concentration of 5×10^8^ microbubbles/mL using a syringe pump (Harvard Apparatus Pump 11 Elite) operating at a flow rate of 20uL/min. Following detection of microbubbles within the tumor, a flash pulse was administered, and images were acquired in the CEUS mode for 20s on a clinical Epiq Elite device. After images were acquired, the animal was removed from the system and woken with antisedan as soon as possible. Image analysis was performed using the Qlabs software on the Epiq device (Phillips, Bethel WA) to create metrics of peak average intensity and wash in slope for microbubble influx as metrics of tumor perfusion. Imaging and analyses were repeated 72 hours post ablation to determine recovery.

### Flow cytometry

Tumors were incubated in Type I collagenase (5 mg/mL, ThermoFisher) and DNAse I (100ug/mL, Sigma) at 37C for 1 hour followed by mechanical homogenization. The disaggregated tumors were filtered through 100 µm Nitex nylon mesh (Genesee) and applied to a differential centrifugation isolation gradient (Lympholyte-M cell separation media, Cedarlane) for cell isolation.

Mice were bled via tail vein, after which blood placed in cell media to prevent clotting, then was RBC lysed. Samples were stained with viability dye (Live Dead BLUE, ThermoFisher) in 1X PBS, followed by a 15-minute incubation with Fc block (anti-mouse functional grade CD16/32). Surface staining was performed in FACS buffer supplemented with 2% normal mouse serum (Valley Biomedical). Cells were permeabilized with Foxp3/Transcription Factor Staining Buffer Set (ThermoFisher). The following antibodies were used across flow cytometry studies described in the results section: CD45 BUV395 (30-F11, BD Biosciences), CD45 FITC (30-F11, ThermoFisher), CD3 BUV563 (145-2C11, BD Biosciences), CD3 APC (145-2C11, BioLegend), CD4 eFluor450 (RM4-5, ThermoFisher), CD8a BUV805 (53-6.7, ThermoFisher), CD11b APC-eFluor780 (M1/70, ThermoFisher), CD11b APC Fire 750 (M1/70, BioLegend), CD19 BV650 (1D3, BD Biosciences), CD19 APC Fire 750 (6D5, BioLegend), CD44 FITC (NIM-R8, BioLegend), Foxp3 PE-Cy5 (236A/E7, ThermoFisher), PD1 BV605 (29F.1A12, BioLegend). Cells were then fixed with 1X FACS Lysis Solution (349202, BD Biosciences).

Samples were acquired on 5-laser Aurora spectral flow cytometer (Cytek) and data were analyzed with FCS Express (De Novo Software). A representative gating strategy for circulating and intratumoral T cells is provided in supplementary Figure 1.

### Survival criteria

The following endpoints were employed for survival studies: euthanasia following tumor outgrowth exceeding 18 mm diameter in any dimension, euthanasia due to weight loss or moribund appearance, or spontaneous death. The following endpoints were censored from survival data: euthanasia due to tumor ulceration.

### Treatment effect assessments

To determine whether the relationships between combined treatments were additive, subtractive, synergistic, or antagonistic, we first created an “effect” metric by calculating the percent reduction in tumor size at day 28 compared to the control group average. We then used two methods were used to determine combination treatment group interactivity: the response additivity model and the Bliss independence model. The response additivity model assumes a synergistic effect when a combination of two therapies yields a greater response than the sum of the individual therapies’ effects:

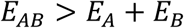

The Bliss independence model assumes a synergistic effect when the two therapies act on different sites of action and when the effect of both therapies is greater than the difference between their sum and product:

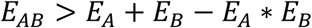

If the combined effect of two therapies exceeded *E*_*AB*_ under both models, we determined that the combined effect was synergistic. If the effect was positive but equal to or between the *E*_*AB*_ values under both models, we determined that the combined effect was additive.

### Statistical Analyses

All results are reported as the mean ± the standard error of the mean (SEM). Percent metabolically active tumor area, immune cell counts and percentages, and area under the curve data comparing two groups were analyzed using an unpaired T-test with Welch’s correction to assess significance. All other area under the curve data and T cell reconstitution data were analyzed with a two-way ANOVA and Tukey post-hoc test to assess significance. Survival data were analyzed using a Kaplan Meier analysis and log-rank Mantel test to assess significance. Rechallenge rejection data were analyzed using a Fisher’s exact test and a multiple comparison post-hoc test to assess significance. Longitudinal tumor growth data were analyzed with a mixed-effects model and Bonferroni correction to assess significance. To identify outliers within experimental groups, the ROUT method was performed where Q = 0.1. To calculate integrated tumor burden, we calculated AUC from day 6 or 7 to day 28 using the trapezoid rule. Statistical significance was assessed at p < 0.05 for all experiments and was calculated using GraphPad Prism 9 (San Diego, USA), with the exception of longitudinal growth control, where significance was calculated using RStudio. Statistical tests are reported in Figure Legends.

## Acknowledgments

We thank Dr. Frederic Padilla for the use of his group’s focused ultrasound system, referred to as System 1 above. We thank the Flow Cytometry Core Facility of the University of Virginia for the use of their cytometers. This core is supported by the National Cancer Institute P30-CA044579 Center Grant.

## Funding

Supported by NIH R01EB030744, R01EB030409, R21CA286367 & R01CA279134 to RJP, the Focused Ultrasound Foundation to RJP, and the University of Virginia’s Focused Ultrasound Cancer Immunotherapy Center to RJP. MRS was supported by an NSF Graduate Research Fellowship and an NIH Cancer Training Grant (T32CA009109).

## Author Contributions

Conceptualization: MRS, RJP, TNJB, MRD Methodology: MRS, LEK, MRD, JRL Formal Analysis: MRS, NZA

Investigation: MRS, NZA

Data Curation: CMR, JVN, DRB, PMD Writing - Original Draft: MRS

Writing - Review and Editing: MRS, NZA, LEK, MRD, TNJB, RJP, HRL Resources: JRL

Funding acquisition: RJP Supervision: RJP

## Notes

### Competing Interest Statement

The authors have declared no competing interest.

## References

1. Knavel, E. M. & Brace, C. L. Tumor ablation: Common modalities and general practices. Techniques in Vascular and Interventional Radiology vol. 16 192–200 Preprint at 10.1053/j.tvir.2013.08.002 (2013).

2. Hatzfeld-Charbonnier, A. S. et al. Influence of heat stress on human monocyte-derived dendritic cell functions with immunotherapeutic potential for antitumor vaccines. J Leukoc Biol 81, 1179 (2007).

3. Mace, T. A., Zhong, L., Kokolus, K. M. & Repasky, E. A. Effector CD8+ T cell IFN-γ production and cytotoxicity are enhanced by mild hyperthermia. International Journal of Hyperthermia 28, 9 (2012).

4. Yu, M. et al. Microwave ablation of primary breast cancer inhibits metastatic progression in model mice via activation of natural killer cells. Cell Mol Immunol 18, 2153 (2020).

5. Xiao, W. et al. The CXCL10/CXCR3 Pathway Contributes to the Synergy of Thermal Ablation and PD-1 Blockade Therapy against Tumors. Cancers (Basel) 15, 1427 (2023).

6. Lemdani, K. et al. Therapeutic and cytotoxic responses after radiofrequency ablation combined to in situ immunomodulation and PD1 blockade in colorectal cancer. Journal of Clinical Oncology 36, e15562–e15562 (2018).

7. Wu, Y. et al. Cryoablation reshapes the immune microenvironment in the distal tumor and enhances the anti-tumor immunity. Front Immunol 13, 930461 (2022).

8. Domingo-Musibay, E., et al. Endogenous Heat-Shock Protein Induction with or Without Radiofrequency Ablation or Cryoablation in Patients with Stage IV Melanoma. Oncologist 22, 1026–e93 (2017).

9. Guo, X. et al. Immunological effect of irreversible electroporation on hepatocellular carcinoma. BMC Cancer 21, (2021).

10. Nuccitelli, R. et al. Nanoelectroablation of Murine Tumors Triggers a CD8-Dependent Inhibition of Secondary Tumor Growth. PLoS One 10, (2015).

11. van den Bijgaart, R. J. E., et al. Thermal and mechanical high-intensity focused ultrasound: perspectives on tumor ablation, immune effects and combination strategies. Cancer Immunol Immunother 66, 247 (2016).

12. Zhang, Y., Deng, J., Feng, J. & Wu, F. Enhancement of antitumor vaccine in ablated hepatocellular carcinoma by high-intensity focused ultrasound. World Journal of Gastroenterology: WJG 16, 3584 (2010).

13. Xia, J. Z. et al. High-intensity focused ultrasound tumor ablation activates autologous tumor-specific cytotoxic T lymphocytes. Ultrasound Med Biol 38, 1363–1371 (2012).

14. Hu, Z. et al. Release of endogenous danger signals from HIFU-treated tumor cells and their stimulatory effects on APCs. Biochem Biophys Res Commun 335, 124 (2005).

15. Kruse, D. E., Mackanos, M. A., O’Connell-Rodwell, C. E., Contag, C. H. & Ferrara, K. W. Short-duration Focused Ultrasound Stimulation of Hsp70 Expression In Vivo. Phys Med Biol 53, 3641 (2008).

16. De Maio, A., Alfieri, G., Mattone, M., Ghanouni, P. & Napoli, A. High-Intensity Focused Ultrasound Surgery for Tumor Ablation: A Review of Current Applications. Radiol Imaging Cancer 6, (2024).

17. Do, H. D. et al. Combination of thermal ablation by focused ultrasound, pFAR4-IL-12 transfection and lipidic adjuvant provide a distal immune response. Explor Target Antitumor Ther 3, 398 (2022).

18. Yang, R. et al. Effects of high-intensity focused ultrasound in the treatment of experimental neuroblastoma. J Pediatr Surg 27, 246–251 (1992).

19. Sheybani, N. D. et al. Combination of thermally ablative focused ultrasound with gemcitabine controls breast cancer via adaptive immunity. J Immunother Cancer 8, e001008 (2020).

20. Hundt, W., O’Connell-Rodwell, C. E., Bednarski, M. D., Steinbach, S. & Guccione, S. In Vitro Effect of Focused Ultrasound or Thermal Stress on HSP70 Expression and Cell Viability in Three Tumor Cell Lines. Acad Radiol 14, 859–870 (2007).

21. Wu, F. et al. Expression of tumor antigens and heat-shock protein 70 in breast cancer cells after high-intensity focused ultrasound ablation. Ann Surg Oncol 14, 1237–1242 (2007).

22. Keum, H. et al. Tissue Ablation: Applications and Perspectives. Advanced Materials 36, 2310856 (2024).

23. Wetterwald, L. et al. Abscopal effect induced by cryoablation in a 55-year-old patient with metastatic dedifferentiated liposarcoma: a case report. Ann Transl Med 12, 94–94 (2024).

24. Bäcklund, M. & Freedman, J. Microwave Ablation and Immune Activation in the Treatment of Recurrent Colorectal Lung Metastases: A Case Report. Case Rep Oncol 10, 383–387 (2017).

25. Thomas, R., Al-Khadairi, G. & Decock, J. Immune Checkpoint Inhibitors in Triple Negative Breast Cancer Treatment: Promising Future Prospects. Front Oncol 10, 600573 (2021).

26. Huang, A. C. & Zappasodi, R. A decade of checkpoint blockade immunotherapy in melanoma: understanding the molecular basis for immune sensitivity and resistance. Nature Immunology 2022 23:5 23, 660–670 (2022).

27. Ju, Y. et al. Barriers and opportunities in pancreatic cancer immunotherapy. npj Precision Oncology 2024 8:1 8, 1–18 (2024).

28. Den Brok, M. H. M. G. M., et al. In Situ Tumor Ablation Creates an Antigen Source for the Generation of Antitumor Immunity. Cancer Res 64, 4024–4029 (2004).

29. Lyu, N. et al. Ablation Reboots the Response in Advanced Hepatocellular Carcinoma With Stable or Atypical Response During PD-1 Therapy: A Proof-of-Concept Study. Front Oncol 10, 580241 (2020).

30. Bonnet, B. et al. Thermal Ablation Combined with Immune Checkpoint Blockers: A 10-Year Monocentric Experience. Cancers (Basel) 16, 855 (2024).

31. Mustafa, A. R., Miyasato, D. & Wehrenberg-Klee, E. Synergizing Thermal Ablation Modalities with Immunotherapy: Enough to Induce Systemic Antitumoral Immunity? Journal of Vascular and Interventional Radiology 35, 185–197 (2024).

32. Chauhan, V. P. et al. Angiotensin inhibition enhances drug delivery and potentiates chemotherapy by decompressing tumour blood vessels. Nat Commun 4, (2013).

33. Matuszewska, K., Pereira, M., Petrik, D., Lawler, J. & Petrik, J. Normalizing tumor vasculature to reduce hypoxia, enhance perfusion, and optimize therapy uptake. Cancers (Basel) 13, 1–19 (2021).

34. Zhu, H. et al. Recombinant human endostatin enhances the radioresponse in esophageal squamous cell carcinoma by normalizing tumor vasculature and reducing hypoxia. Scientific Reports 2015 5:1 5, 1–9 (2015).

35. Huang, H. et al. VEGF suppresses T-lymphocyte infiltration in the tumor microenvironment through inhibition of NF-κB-induced endothelial activation. FASEB Journal 29, 227–238 (2015).

36. Dirkx, A. E. M. et al. Anti-angiogenesis therapy can overcome endothelial cell anergy and promote leukocyte-endothelium interactions and infiltration in tumors. The FASEB Journal 20, 621–630 (2006).

37. McDonald, D. M. & Baluk, P. Imaging of angiogenesis in inflamed airways and tumors: newly formed blood vessels are not alike and may be wildly abnormal: Parker B. Francis lecture. Chest 128, 602S–608S (2005).

38. Fluctuations in red cell flux in tumor microvessels can lead to transient hypoxia and reoxygenation in tumor parenchyma - PubMed. https://pubmed.ncbi.nlm.nih.gov/8968110/.

39. Piali, L., Fichtd, A., Terpe, H. J., Imhof, B. A. & Gisler, R. H. Endothelial vascular cell adhesion molecule 1 expression is suppressed by melanoma and carcinoma. Journal of Experimental Medicine 181, 811–816 (1995).

40. Griffioen, A. W., Damen, C. A., Martinotti, S., Blijham, G. H. & Groenewegen, G. Endothelial intercellular adhesion molecule-1 expression is suppressed in human malignancies: The role of angiogenic factors. Cancer Res 56, 1111–1117 (1996).

41. Griffioen, A. W., Damen, C. A., Blijham, G. H. & Groenewegen, G. Tumor Angiogenesis Is Accompanied by a Decreased Inflammatory Response of Tumor-Associated Endothelium. Blood 88, 667–673 (1996).

42. Tromp, S. C. et al. Tumor angiogenesis factors reduce leukocyte adhesion in vivo. Int Immunol 12, 671–676 (2000).

43. Dirkx, A. E. M. et al. Anti-angiogenesis therapy can overcome endothelial cell anergy and promote leukocyte-endothelium interactions and infiltration in tumors. The FASEB Journal 20, 621–630 (2006).

44. Muller, W. A. Leukocyte-Endothelial Cell Interactions in the Inflammatory Response. Laboratory Investigation 2002 82:5 82, 521–534 (2002).

45. Farsaci, B. et al. Immune consequences of decreasing tumor vasculature with antiangiogenic tyrosine kinase inhibitors in combination with therapeutic vaccines. Cancer Immunol Res 2, 1090–1102 (2014).

46. Draghiciu, O., Nijman, H. W., Hoogeboom, B. N., Meijerhof, T. & Daemen, T. Sunitinib depletes myeloid-derived suppressor cells and synergizes with a cancer vaccine to enhance antigen-specific immune responses and tumor eradication. Oncoimmunology 4, 1–11 (2015).

47. Huang, Y. et al. Vascular normalizing doses of antiangiogenic treatment reprogram the immunosuppressive tumor microenvironment and enhance immunotherapy. Proc Natl Acad Sci U S A 109, 17561–17566 (2012).

48. Shrimali, R. K. et al. Antiangiogenic agents can increase lymphocyte infiltration into tumor and enhance the effectiveness of adoptive immunotherapy of cancer. Cancer Res 70, 6171 (2010).

49. Shi, S. et al. Combining Antiangiogenic Therapy with Adoptive Cell Immunotherapy Exerts Better Antitumor Effects in Non-Small Cell Lung Cancer Models. PLoS One 8, (2013).

50. Tao, L., Huang, G., Shi, S. & Chen, L. Bevacizumab improves the antitumor efficacy of adoptive cytokine-induced killer cells therapy in non-small cell lung cancer models. Medical Oncology 31, 1–8 (2014).

51. E. A., et al. Combined antiangiogenic and anti-PD-L1 therapy stimulates tumor immunity through HEV formation. Sci Transl Med 9, (2017).

52. Schmittnaegel, M. et al. Dual angiopoietin-2 and VEGFA inhibition elicits antitumor immunity that is enhanced by PD-1 checkpoint blockade. Sci Transl Med 9, (2017).

53. Li, Q. et al. Low-dose anti-angiogenic therapy sensitizes breast cancer to PD-1 blockade. Clinical Cancer Research 26, 1712–1724 (2020).

54. Yasuda, S. et al. Simultaneous blockade of programmed death 1 and vascular endothelial growth factor receptor 2 (VEGFR2) induces synergistic anti-tumour effect in vivo. Clin Exp Immunol 172, 500–506 (2013).

55. Li, Y. et al. Treatment with a VEGFR-2 antibody results in intra-tumor immune modulation and enhances anti-tumor efficacy of PD-L1 blockade in syngeneic murine tumor models. PLoS One 17, 1–16 (2022).

56. Meder, L. et al. Combined VEGF and PD-L1 blockade displays synergistic treatment effects in an autochthonous mouse model of small cell lung cancer. Cancer Res 78, 4270–4281 (2018).

57. Di Tacchio, M. et al. Tumor vessel normalization, immunostimulatory reprogramming, and improved survival in glioblastoma with combined inhibition of PD-1, angiopoietin-2, and VEGF. Cancer Immunol Res 7, 1910–1927 (2019).

58. Kohei Shigeta 1, 2,#, Meenal Datta1,#, Tai Hato1, 3. Dual PD-1 and VEGFR-2 blockade promotes vascular normalization and enhances anti-tumor immune responses in HCC. Hepatology 176, 139–148 (2020).

59. Goda, N. et al. The ratio of CD8 + lymphocytes to tumor-infiltrating suppressive FOXP3 + effector regulatory T cells is associated with treatment response in invasive breast cancer. *Discover*. Oncology 13, 27 (2022).

60. Preston, C. C. et al. The Ratios of CD8+ T Cells to CD4+CD25+ FOXP3+ and FOXP3-T Cells Correlate with Poor Clinical Outcome in Human Serous Ovarian Cancer. PLoS One 8, e80063 (2013).

61. Foucquier, J. & Guedj, M. Analysis of drug combinations: current methodological landscape. Pharmacol Res Perspect 3, e00149 (2015).

62. Hobbs, S. J. & Nolz, J. C. Targeted Expansion of Tissue-Resident CD8+ T Cells to Boost Cellular Immunity in the Skin. Cell Rep 29, 2990 (2019).

63. Clark, R. A. et al. Skin effector memory T cells do not recirculate and provide immune protection in alemtuzumab-treated CTCL patients. Sci Transl Med 4, 117ra7 (2012).

64. Goplen, N. P., et al. Tissue-resident CD8+ T cells drive age-associated chronic lung sequelae after viral pneumonia. Sci Immunol 5, (2020).

65. Gao, M. et al. Therapy With Carboplatin and Anti-PD-1 Antibodies Before Surgery Demonstrates Sustainable Anti-Tumor Effects for Secondary Cancers in Mice With Triple-Negative Breast Cancer. Front Immunol 11, 1–16 (2020).

66. Grosso, J. F. & Jure-Kunkel, M. N. CTLA-4 blockade in tumor models: an overview of preclinical and translational research. Cancer Immun 13, 5 (2013).

67. Jin, Y. et al. Different syngeneic tumors show distinctive intrinsic tumor-immunity and mechanisms of actions (MOA) of anti-PD-1 treatment. Sci Rep 12, 1–18 (2022).

68. Debien, V. et al. Immunotherapy in breast cancer: an overview of current strategies and perspectives. NPJ Breast Cancer 9, 1–10 (2023).

69. Fite, B. Z. et al. Immune modulation resulting from MR-guided high intensity focused ultrasound in a model of murine breast cancer. Sci Rep 11, 927 (2021).

70. Silvestrini, M. T., et al. Priming is key to effective incorporation of image-guided thermal ablation into immunotherapy protocols. JCI Insight 2, e90521 (2017).

71. Han, X. et al. In situ thermal ablation of tumors in combination with nano-adjuvant and immune checkpoint blockade to inhibit cancer metastasis and recurrence. Biomaterials 224, (2019).

72. Chavez, M. et al. Distinct immune signatures in directly treated and distant tumors result from TLR adjuvants and focal ablation. Theranostics 8, 3611–3628 (2018).

73. Yang, X. et al. High-intensity focused ultrasound ablation combined with immunotherapy for treating liver metastases: A prospective non-randomized trial. PLoS One 19, e0306595 (2024).

74. High-intensity focused ultrasound thermal ablation boosts the efficacy of immune checkpoint inhibitors in advanced cancers with liver metastases: A single-center retrospective cohort study. https://www.spandidos-publications.com/10.3892/ol.2025.14871.

75. Tang, R. et al. Novel combination strategy of high intensity focused ultrasound (HIFU) and checkpoint blockade boosted by bioinspired and oxygen-supplied nanoprobe for multimodal imaging-guided cancer therapy. J Immunother Cancer 11, e006226 (2023).

76. Ran, L. F. et al. T-lymphocytes from focused ultrasound ablation subsequently mediate cellular antitumor immunity after adoptive cell transfer immunotherapy. Front Immunol 14, 1155229 (2023).

77. Ran, L. F. et al. Specific antitumour immunity of HIFU-activated cytotoxic T lymphocytes after adoptive transfusion in tumour-bearing mice. International Journal of Hyperthermia 32, 204–210 (2016).

78. Ashar, H. et al. Enabling Chemo-Immunotherapy with HIFU in Canine Cancer Patients. Ann Biomed Eng 52, 1859–1872 (2024).

79. Chiu, L. C., Wu, S. K., Lin, W. L. & Chen, G. S. Synergistic Effects of Nanodrug, Ultrasound Hyperthermia, and Thermal Ablation on Solid Tumors-An Animal Study. IEEE Trans Biomed Eng 64, 2880–2889 (2017).

80. Wong, A. W. et al. Ultrasound ablation enhances drug accumulation and survival in mammary carcinoma models. J Clin Invest 126, 99 (2015).

81. Su, S., Wang, Y., Lo, E. M., Tamukong, P. & Kim, H. L. High-intensity focused ultrasound ablation to increase tumor-specific lymphocytes in prostate cancer. Transl Oncol 53, 102293 (2025).

82. Liao, Y. et al. High-intensity focused ultrasound thermal ablation boosts the efficacy of immune checkpoint inhibitors in advanced cancers with liver metastases: A single-center retrospective cohort study. Oncol Lett 29, 124 (2025).

83. Liu, F. et al. Boosting high-intensity focused ultrasound-induced anti-tumor immunity using a sparse-scan strategy that can more effectively promote dendritic cell maturation. J Transl Med 8, 1–12 (2010).

84. Thim, E. A. et al. Solid tumor treatment via augmentation of bioactive C6 ceramide levels with thermally ablative focused ultrasound. Drug Deliv Transl Res 13, 3145 (2023).

85. Chavez, M. et al. Distinct immune signatures in directly treated and distant tumors result from TLR adjuvants and focal ablation. Theranostics 8, 3611–3628 (2018).

86. Abe, S. et al. Original research: Combination of ultrasound-based mechanical disruption of tumor with immune checkpoint blockade modifies tumor microenvironment and augments systemic antitumor immunity. J Immunother Cancer 10, 3717 (2022).

87. Thim, E. A. et al. Focused ultrasound ablation of melanoma with boiling histotripsy yields abscopal tumor control and antigen-dependent dendritic cell activation. Theranostics 14, 1647 (2024).

