## Supplemental Materials for "VEGFR2 blockade converts thermally ablative focused ultrasound into a potent driver of T cell-dependent anti-tumor immunity"

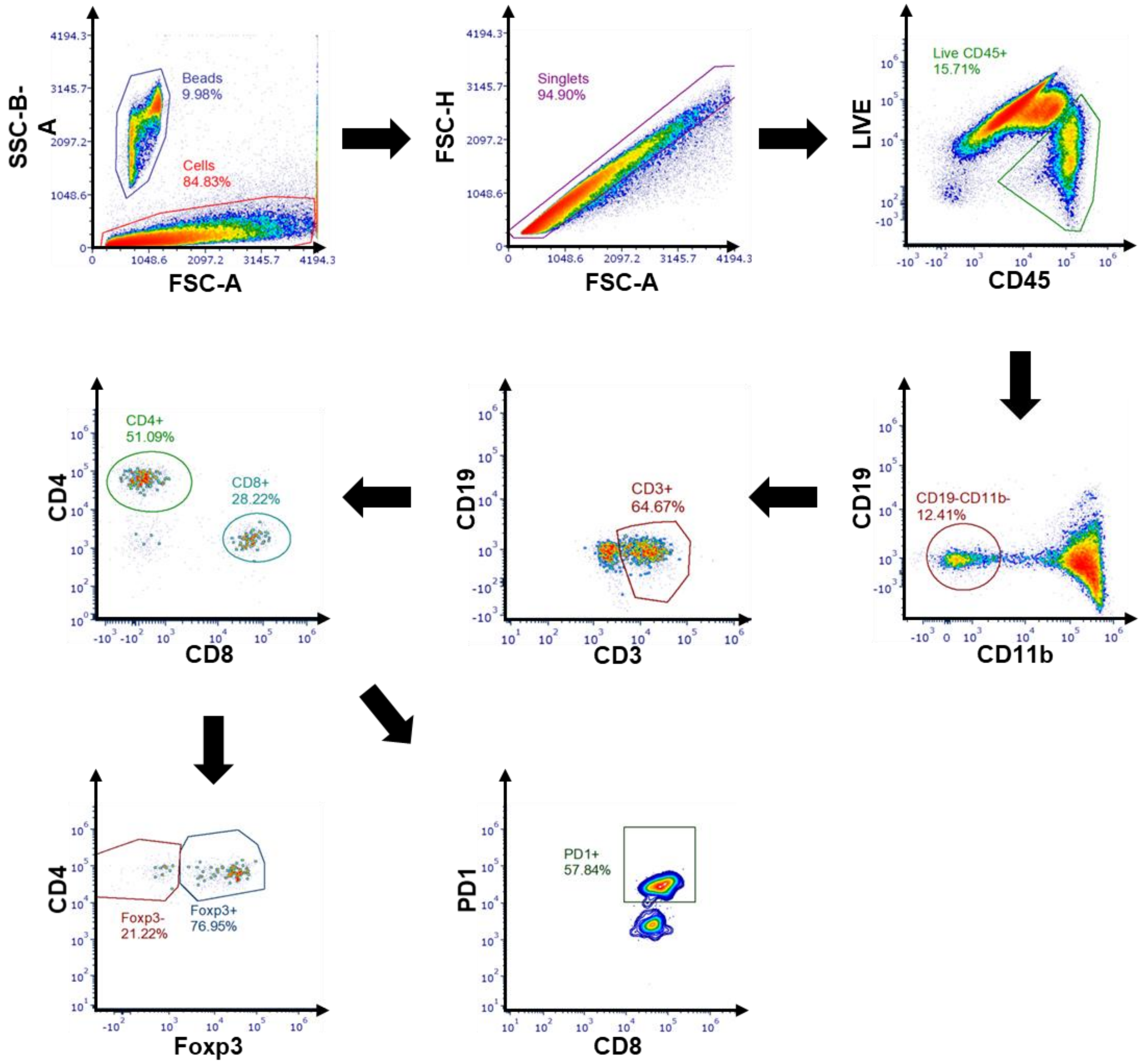


**Figure S1.** Intratumoral immune staining gating strategy post-DC101 treatment.

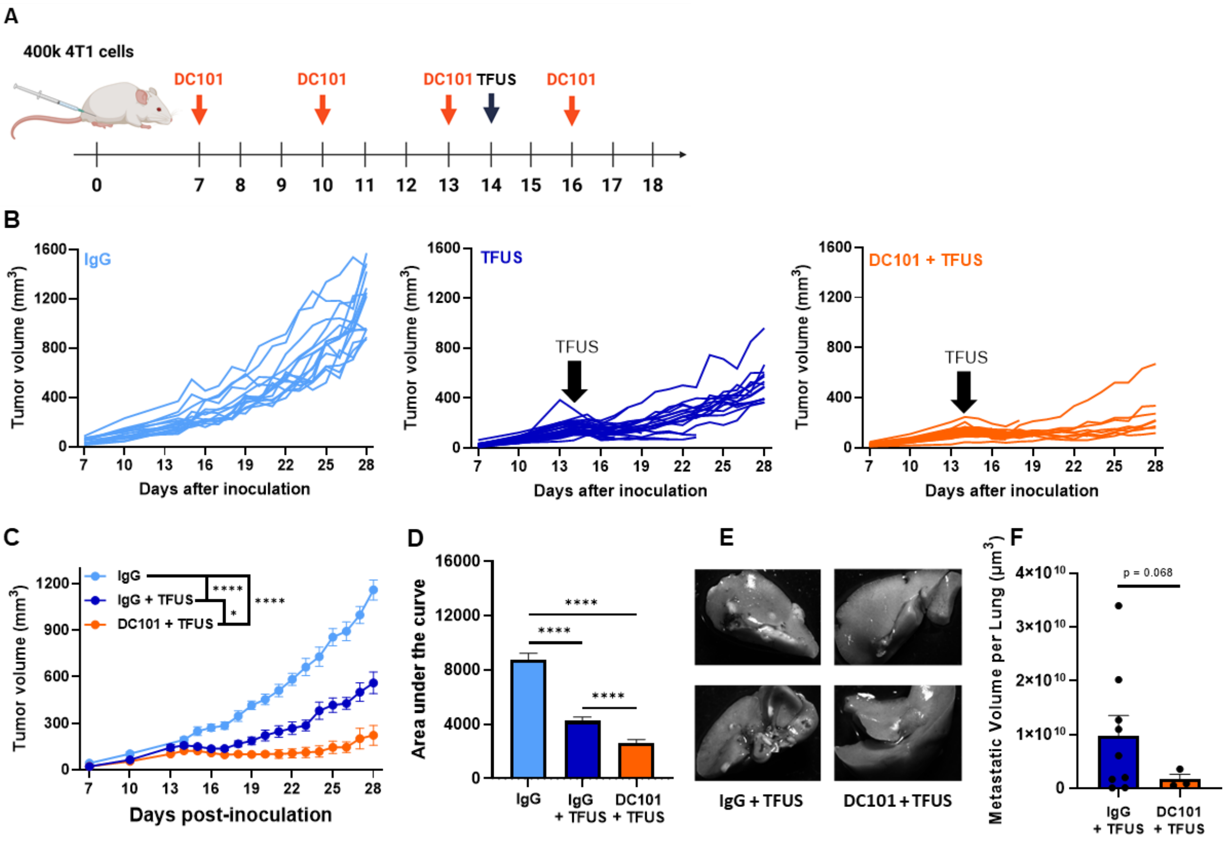


**Figure S2. DC101 and TFUS curb primary tumor growth and prevent distant metastases of 4T1 tumors.** A) Timeline for inoculation and treatment. B) Individual growth curves. C) Grouped tumor growth over time. Two-way, repeated measures ANOVA. ****P < 0.0001, *P = 0.0219. D) Area under the curve (AUC) for tumor growth. One-way ANOVA with Tukey's test. ****P < 0.0001. E) Representative lungs stained with India ink, with metastases visible in white. F) Estimated metastatic lung volume quantified in ImageJ. Welch's t-test.

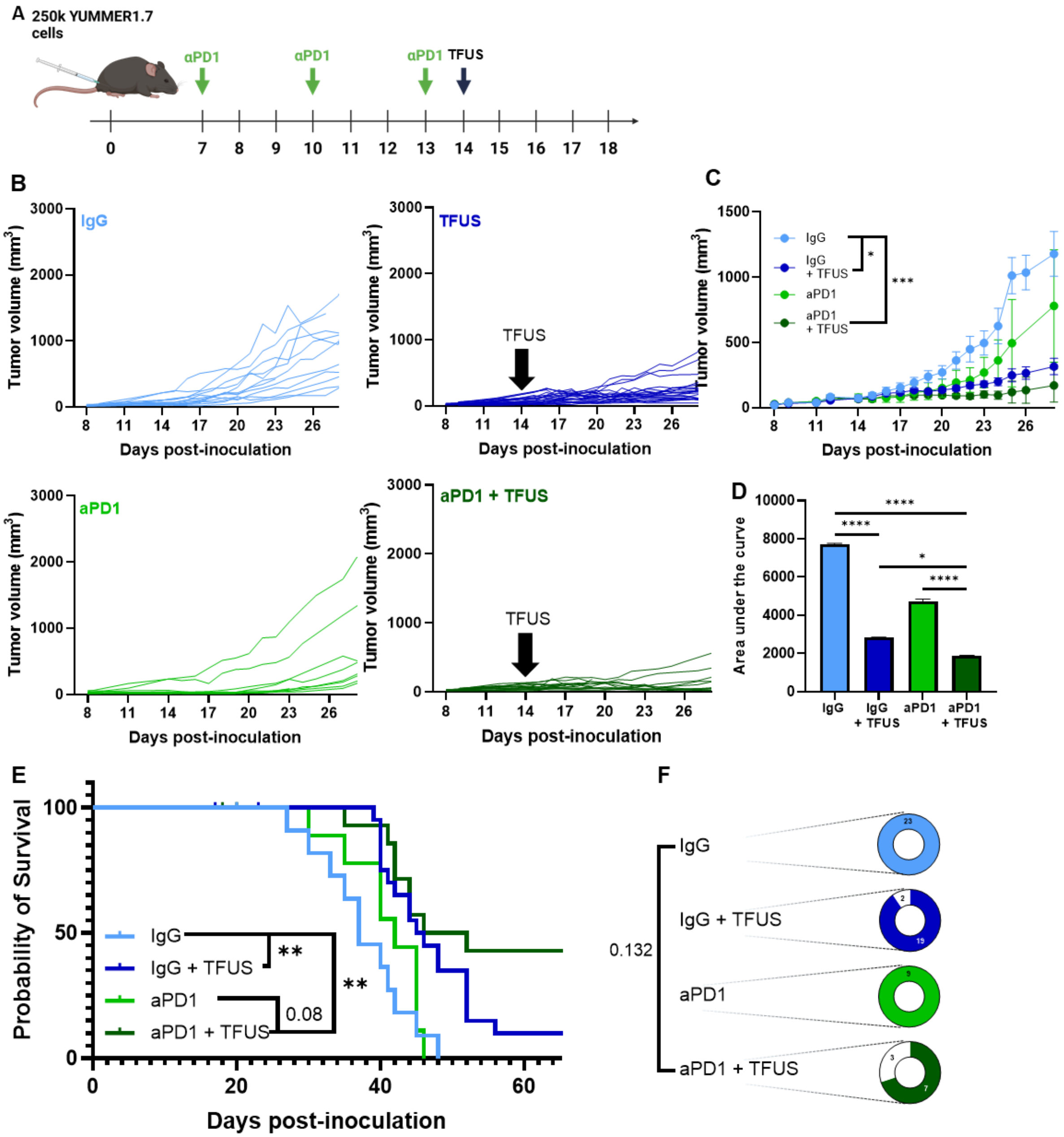


**Figure S3. TFUS and aPD1 cooperate to control tumor growth in YUMMER1.7 tumors.** A) Timeline for inoculation and treatment. B) Individual growth curves. C) Grouped tumor growth over time. Two-way, repeated measures ANOVA. *P = 0.0280, ***P = 0.0009. D) Area under the curve (AUC) for tumor growth. Two-way ANOVA with Tukey's test. ****P < 0.0001, *P = 0.0373. E) Kaplan-Meier curve depicting overall survival. Mantel-Cox log-rank test. **P < 0.01. F) Tumor eradication fraction (white shading). Fisher's exact test.
